# Mapping the bioimaging marker of Alzheimer’s disease based on pupillary light response-driven brain-wide fMRI in awake mice

**DOI:** 10.1101/2023.12.20.572613

**Authors:** Xiaochen Liu, David Hike, Sangcheon Choi, Weitao Man, Chongzhao Ran, Xiaoqing Alice Zhou, Yuanyuan Jiang, Xin Yu

## Abstract

Pupil dynamics has emerged as a critical non-invasive indicator of brain state changes. In particular, pupillary-light-responses (PLR) in Alzheimer’s disease (AD) patients may be used as biomarkers of brain degeneration. To characterize AD-specific PLR and its underlying neuromodulatory sources, we combined high-resolution awake mouse fMRI with real-time pupillometry to map brain-wide event-related correlation patterns based on illumination-driven pupil constriction (*P_c_*) and post-illumination pupil dilation recovery (amplitude, *P_d_*, and time, *T*). The *P_c_*-driven differential analysis revealed altered visual signal processing coupled with reduced thalamocortical activation in AD mice compared with the wild-type normal mice. In contrast, the post-illumination pupil dilation recovery-based fMRI highlighted multiple brain areas related to AD brain degeneration, including the cingulate cortex, hippocampus, septal area of the basal forebrain, medial raphe nucleus, and pontine reticular nuclei (PRN). Also, brain-wide functional connectivity analysis highlighted the most significant changes in PRN of AD mice, which serves as the major subcortical relay nuclei underlying oculomotor function. This work combined non-invasive pupil-fMRI measurements in preclinical models to identify pupillary biomarkers based on neuromodulatory dysfunction coupled with AD brain degeneration.

## Introduction

Pupil diameter changes reflect cognitive processing^1–4^, presenting a unique non-invasive index of brain state fluctuation in normal and degenerative brains, e.g. Alzheimer’s Disease (AD) ^5–23^. Although several studies associate pupillary responses during cognitive tasks with the risk for AD, it remains highly speculative given the lack of understanding of its mechanistic linkage ^24–28^. Multiple neuronal sources drive the highly varied pupil dynamics, including pupillary light responses (PLR), through autonomic neuromodulation to control pupil dilation and constriction^9–11,29–36^. Since neuromodulatory dysfunction is often observed in degenerative AD brains^37–40^, it is plausible to identify dynamic PLR serving as novel functional biomarkers based on converging effects from malfunctional neuromodulatory pathways of AD brains. The ongoing challenge is to bridge the complex PLR with the underlying brain-wide functional alteration in degenerative AD brains.

Combining whole-brain fMRI with real-time pupillometry provides a non-invasive mapping scheme to identify pupil-fMRI relationships across different species. Existing human fMRI studies with pupillometry have revealed pupil dynamic changes associated with the activation of subcortical brain nuclei along the ascending arousal network, as well as the salience network including the cingulate cortex and insula^32,41–49^. Interestingly, opposite arousal correlation features between eye movements and resting-state fMRI have been revealed between the cortex and subcortical nuclei in both non-human primates and rodent brains ^50–52^. The highly varied pupil-dependent brain state changes have been investigated by electrophysiological recordings ^53–55^ and fiber photometry with fMRI ^56–58^ in both anesthetized and awake rodent brains. In particular, the cortical noradrenergic and cholinergic axonal activities are coupled with either rapid or long-lasting pupil dilation, respectively,^56^, indicating neuromodulatory specificity to pupil dynamics. Meanwhile, vagus nerve stimulation (VNS) has been reported to induce pupil dilation through evoked basal-forebrain cholinergic axonal activity^59^. Since the afferent vagal nerves project diverse functional nuclei in brainstem, e.g. LC and raphe nuclei, via the nucleus tractus solitarii (NTS)^60–62^, the emerged neuromodulation on pupil dynamics is well exemplified by VNS paradigms. These studies have illustrated the power of combined pupil-fMRI to reveal complex neuromodulatory networks underlying pupil dynamic changes in diseased animal models.

To elucidate the impaired neuromodulatory pathways underlying AD-specific pupillary light response (PLR), we implemented awake mouse fMRI with simultaneous real-time pupillometry. Using an implantable radiofrequency (RF) surface coil, which also served as a headpost for head-fixation during scanning, we acquired robust high-resolution brain-wide BOLD functional maps (100×100×200µm) of awake mice undergoing visual stimulation. The epoch-specific PLR metrics were measured during awake mouse fMRI. These included the illumination-induced pupil constriction amplitude (*P_c_*), and the post-illumination pupil dilation recovery time (T) and amplitude (*P_d_*), Using the time-varied PLR dynamic features, the *P_c_*-based fMRI differential maps between AD and wild-type (WT) mice were presented to identify altered brain activation in the visual pathways. In contrast, post-illumination pupil dilation (*T*, *P_d_*)-based fMRI differential maps highlighted several cortical (e.g. cingulate and retrosplenial cortex (RSP)) and subcortical regions (e.g. septal areas in the basal forebrain, hippocampus, median raphe nucleus (MnR), and pontine reticular nucleus (PRN)) involved in the altered neuromodulation underlying the AD-specific PLR. Meanwhile, ROI-based correlation analysis also identified altered functional connectivity among these PLR-related neuromodulatory nuclei, highlighting the most significant connectivity changes originating from the PRN between AD and WT mice. These results layout a novel pupil-fMRI mapping scheme to identify unique PLR features as a non-invasive biomarker of AD brains, based on underlying functional mapping differences of neuromodulatory circuits.

## Results

### Epoch-specific pupillary light response recording during awake mouse fMRI

The real-time pupillometry was simultaneously performed inside the 14 Tesla scanner with high-resolution echo-planar-imaging (EPI) scans of awake mice (**Fig 1A**). The awake mouse real-time pupillometry setup includes i. MR-compatible miniaturized complementary metal-oxide-semiconductor (CMOS) chips with adjustable lens to keep at least 2 cm distance from the mouse to avoid B0 field distortion, ii. Plane mirror to reflect pupil toward the camera mounted on the animal holder, iii. The optical fibers to deliver the switchable blue/green light pulses (530nm and 490nm, flashed at 5Hz and 5.1Hz respectively) for visual stimulation and red light (660 nm) to illuminate the eye for pupillometry as mice are less sensitive to red light ^63^.

**Figure 1.**
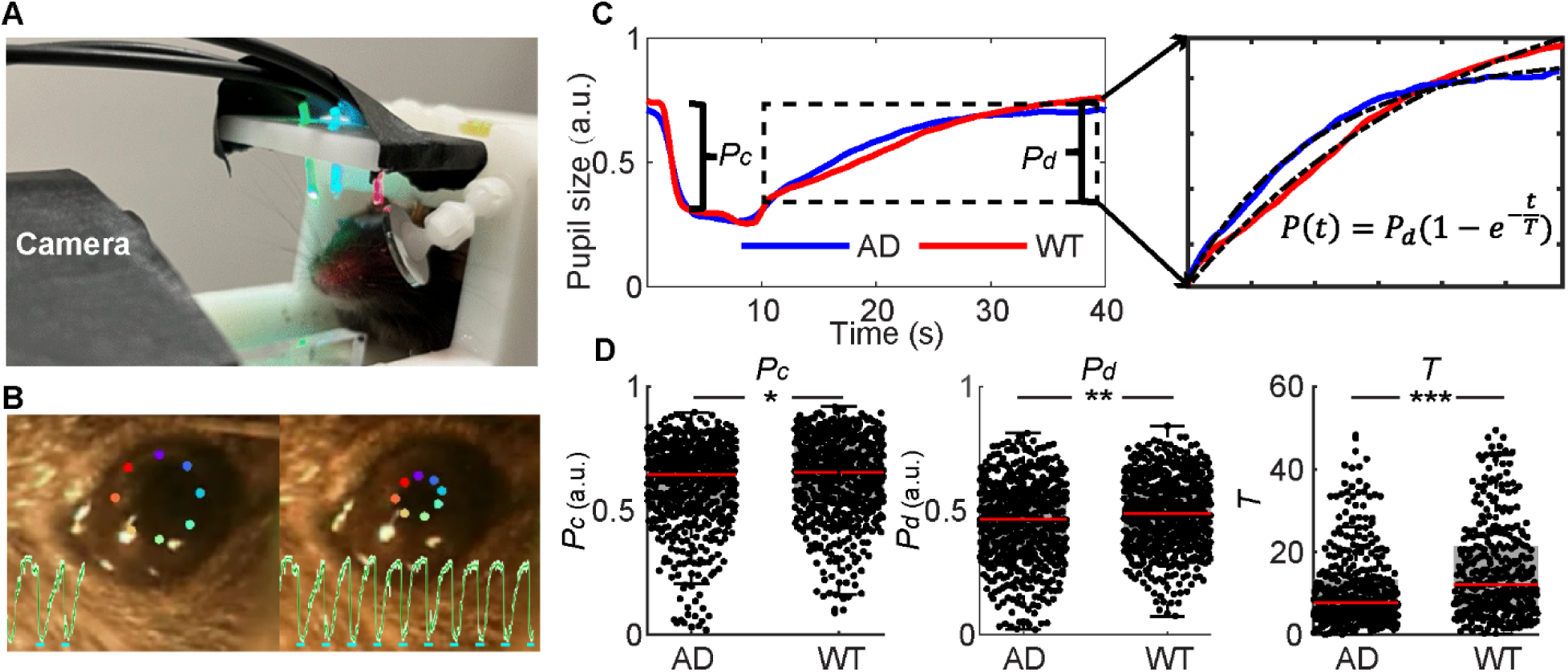
PLR measurements of awake WT and AD mice during fMRI scanning. A) The snapshot of the awake mouse fMRI setup with real-time pupillometry and the visual stimulation setup through optical fibers. B) The representative pupillometry results of an awake mouse with time-varied images of the pupil recorded during fMRI scanning (colored dots define the pupil size based on the DeepLabCut; green trace showed the pupil dynamic changes with visual stimulation in a block design; cyan lines showed stimulation on periods). C) The normalized curves showed averaged pupil light response from AD and WT mice (the bracket indicates the *P_c_*; dotted box highlights the dilation recovery period for exponential fitting function to extract *T* and *P_d_*). D) The PLR feature-specific box plots of *P_c_*, *P_d_*, and *T* between AD and WT mice. (WT: n=9 with 580 stimulation on/off epochs, denoted by black dots, 580; AD: n=9 with 600 stimulation on/off epochs, denoted by black dots; * means p<0.05. ** means p<0.01. *** means p<0.001.)

Mice were implanted with a RF surface coil that served as the headpost for head-fixation while inside the animal holder during scanning (detailed setup is described in ^64^). **Fig 1B** showed the representative trace of PLR from an awake mouse, which was measured by DeepLabCut ^65^. To produce better accuracy of the PLR measurement, we also implemented grayscale-based pupil size measurements to verify different PLR dynamic features across different trials of pupillometry recordings (**Supplementary Fig 1**). The mean PLRs of two groups of mice (AD vs. WT) were plotted in **Fig 1C**, which showed the dynamic PLR features at constriction (i.e. *P_c_*) and dilation recovery phases (*P_d_*, *T*). The recovery time (T) was quantified based on exponential fitting of the pupillary dilation curve from each stimulation on/off epoch. **Fig 1D** showed quantitative comparison of the PLR features, demonstrating significantly reduced pupil size changes in AD mice in terms of the *P_c_* and *P_d_*. It should be noted that the dilation recovery time (*T)* of AD mice presented the most salient changes among the three PLR features. This result suggests that PLR-based fMRI mapping can be used to identify brain-wide functional changes using alterations of AD-driven PLR features acquired during awake mice fMRI.

### Brain-wide BOLD-fMRI of awake AD and WT mice with visual stimulation

High-resolution fMRI was applied to acquire EPI images of awake mice with 100×100×200µm resolution using a horizontal 14T MRI scanner (**Supplementary Fig 2**). During awake mouse fMRI scanning, motion artifacts due to voluntary movement of mice led to distorted images as shown in Supplementary Fig 2E, which can be removed from the 3D time series for regression analysis (**Supplementary Movie 1**). **Fig 2** showed the visual stimulation-evoked brain-wide activation in the visual cortex (VC), superior colliculus (SC), lateral geniculate nucleus (LGN), retrosplenial cortex (RSP), and anterior cingulate cortex (ACA) in both WT and AD mice. Robust positive BOLD signals were detected in the ROI specific time course data of both groups segmented from the Allen mouse brain atlas ^66^ (**Fig 3**), demonstrating the reliable visual stimulation-evoked fMRI signals in awake WT and AD mice.

**Figure 2.**
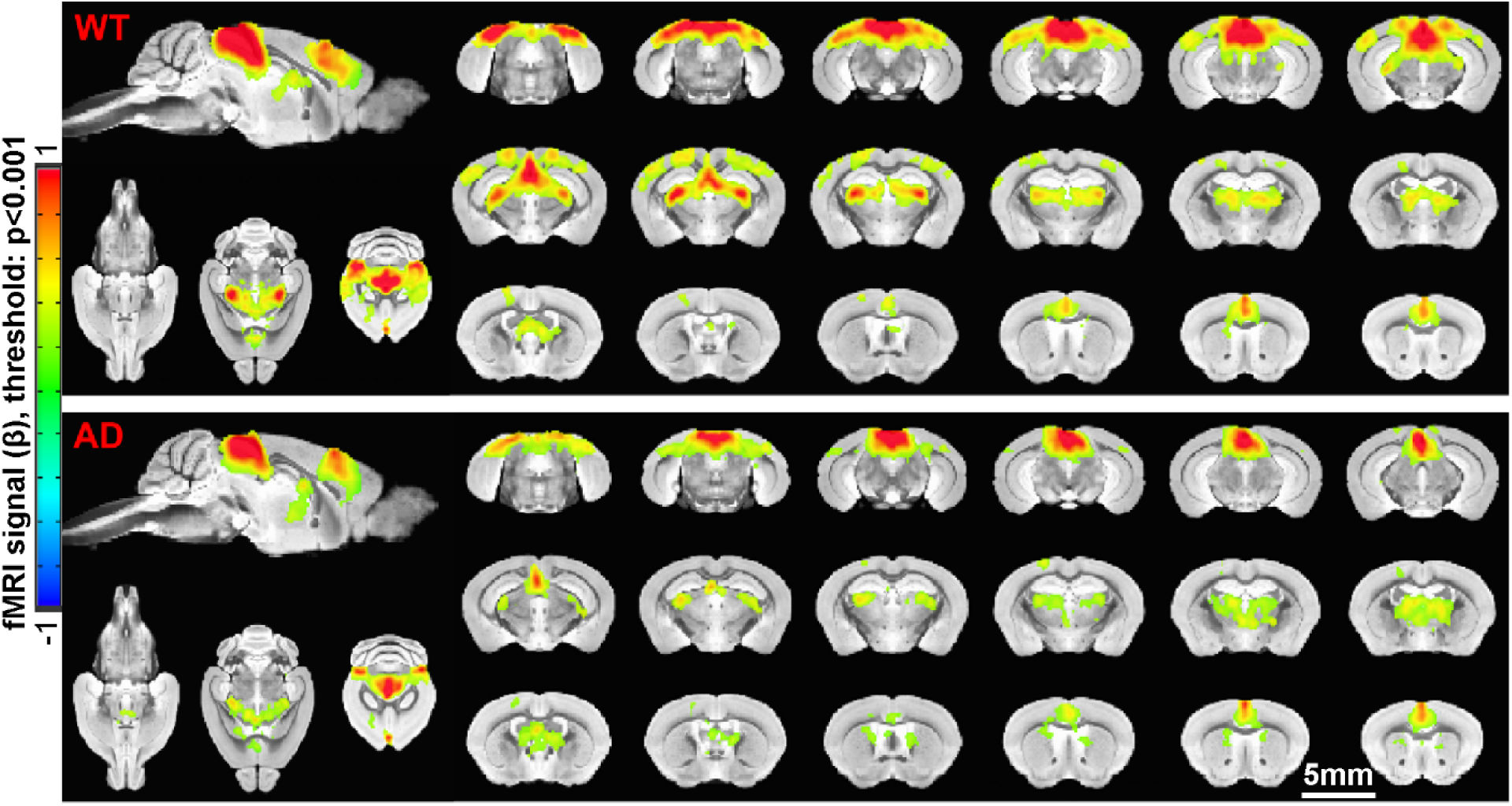
The brain-wide BOLD functional maps of awake WT and AD mice with visual stimulation. The functional maps of awake mice show significant BOLD activation in the visual cortex (VC), superior colliculus (SC), retrosplenial cortex (RSP), lateral geniculate nucleus (LGN), and anterior cingulate area (ACA) from both WT (upper panel) (n=13 mice with 58 trials) and AD (lower panel) mice (n= 9 mice with 60 trials; p<0.001, minimum cluster size=200 voxels).

**Figure 3.**
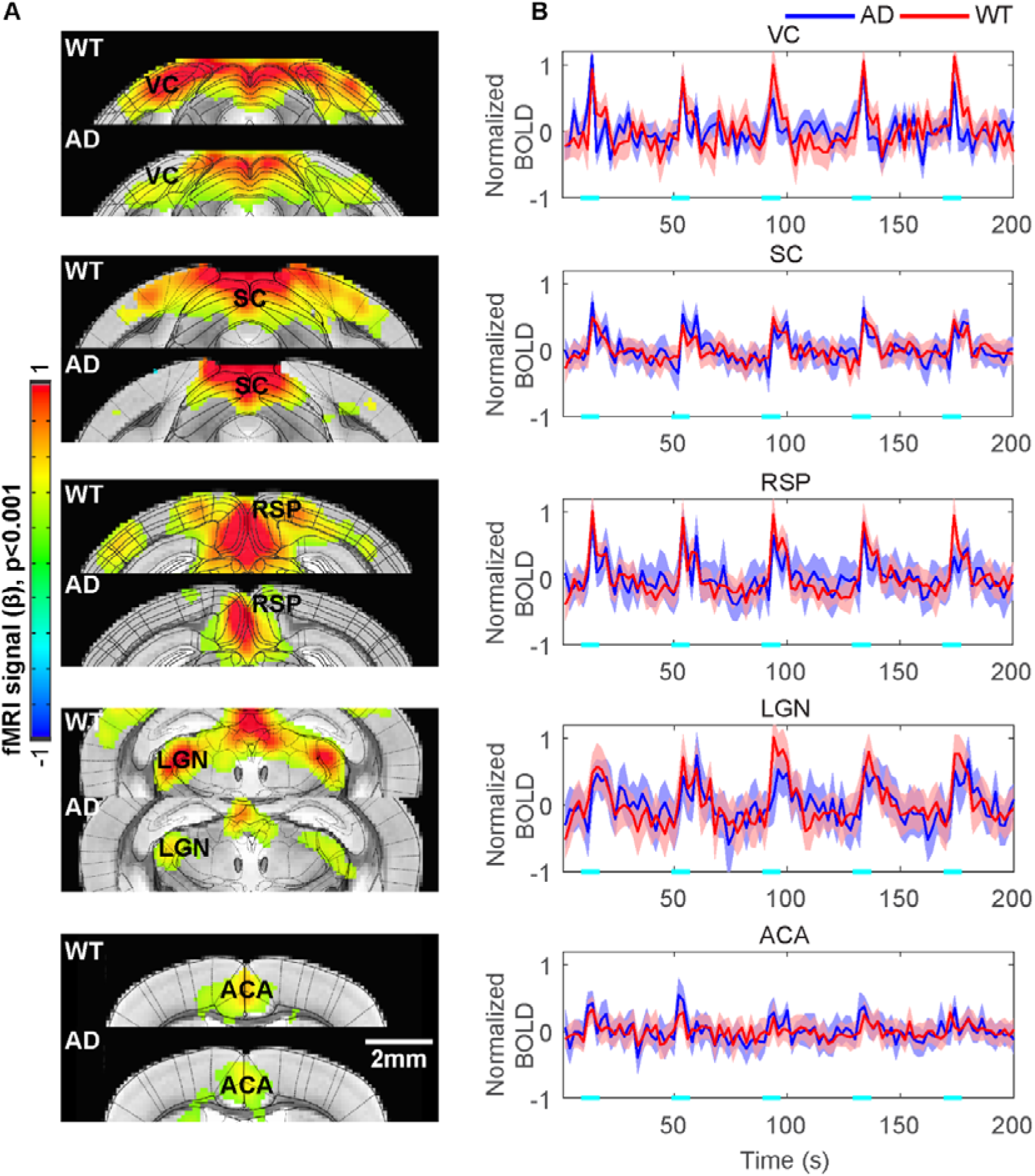
The time courses of visual stimulation evoked BOLD fMRI from ROIs of Allen brain atlas. A) The functional maps overlaid with the brain atlas highlight activated brain regions: visual cortex (VC), superior colliculus (SC), retrosplenial cortex (RSP), lateral geniculate nucleus (LGN), and anterior cingulate area (ACA) from AD and WT mice. B) The averaged time course based on the ROIs from brain atlas, demonstrating the evoked positive BOLD signal changes with the 8s visual stimulation (5Hz 530nm and 5.1Hz 490nm 20ms light pluses). Each graph displays the average of 116 sets of 5 stimulation epochs for 13 WT mice and 120 sets of 5 stimulation epochs for 9 AD mice. Shaded regions represent standard error. Cyan lines represent the 8s stimulation duration (WT: n=13 mice with 58 trials; AD: n=9 mice with 60 trials).

To investigate the differences of functional maps between the two groups, we first integrated the *P_c_*-based amplitude modulated (AM) hemodynamic response function (HRF) as the regressor across different trials. Since the varied pupil constriction at each stimulation on/off epoch altered retinal illumination, specific visual pathway activation would be better quantified by integrating the *P_c_*-based AM regression scheme. Quantitative group analysis was performed to produce voxel-wise differential maps between WT and AD mice. **Fig 4** showed the significantly higher BOLD responses in WT when compared to AD mice in the primary and secondary visual cortices and LGN, but little difference was detected in the SC and cingulate cortex. In particular, well separated higher visual cortical areas, including the adjacent RSP, were highlighted in the differential maps (**Fig 4B**). These results indicate that the higher order thalamocortical processing of visual signals is impaired in the AD mice based on the *P_c_*-fMRI AM regression analysis.

**Figure 4.**
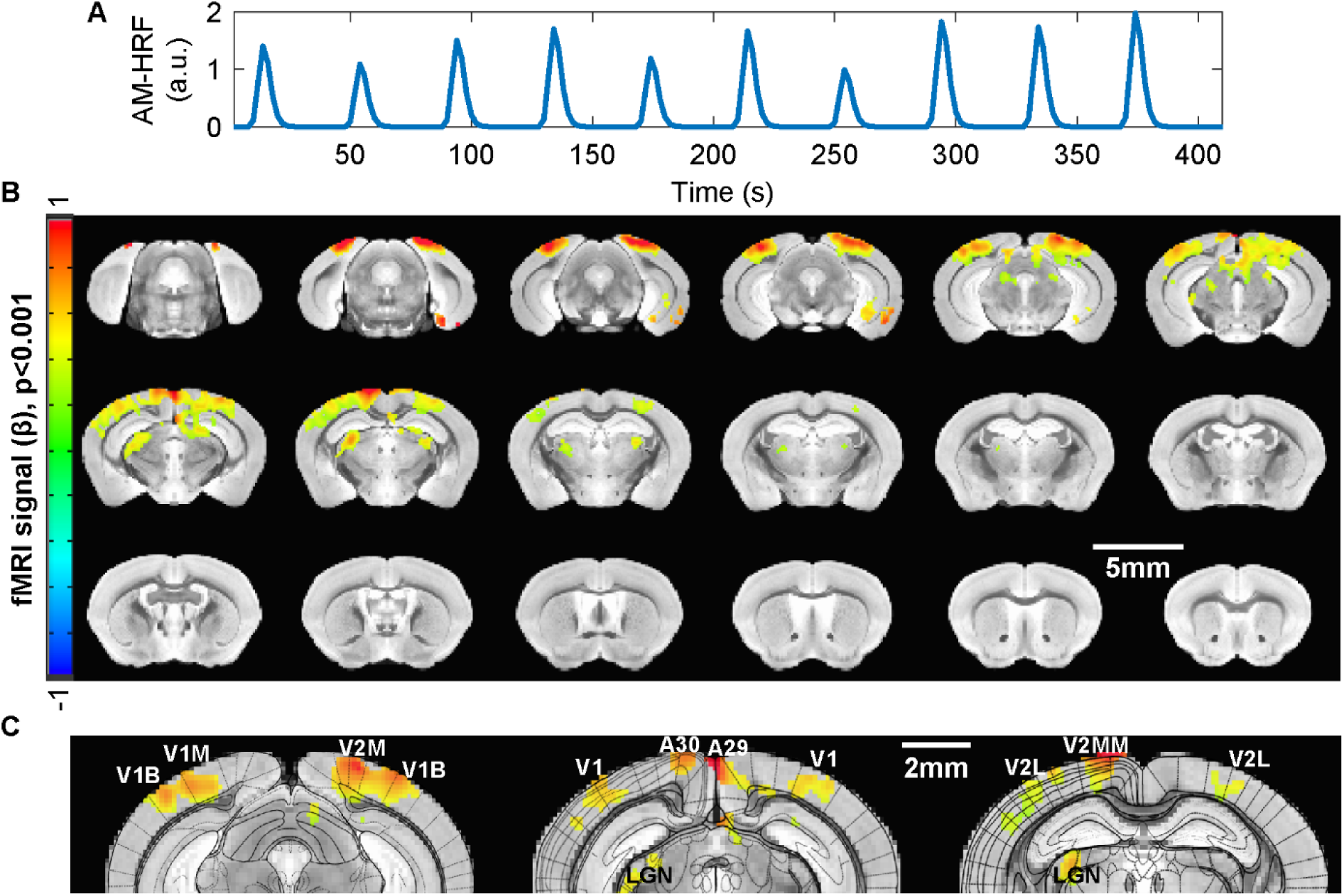
Differential analysis of *P_c_*-based fMRI correlation maps between WT and AD mice. A) A representative regression function derived from the *P_c_*-based amplitude modulation through AM1 function of AFNI. B) The brain-wide *P_c_*-based fMRI differential maps (WT-AD) show the significantly higher BOLD signals detected along visual pathways in WT mice than AD mice (WT: n=9 mice with 20 sections; AD: n=9 mice with 19 sections; p<0.001, minimum cluster size=200 voxels). C) The enlarged functional maps overlaid with the brain atlas highlight the higher order visual pathways: visual cortex (V1, V2), retrosplenial cortex (A30, A29), and lateral geniculate nucleus (LGN).

### Neuromodulation of the post-illumination pupil dilation recovery

Likewise, to the pupil-based brain activation mapping of visual pathways, the potential neuromodulatory nuclei underlying post-illumination pupil dilation were investigated based on the pupil-fMRI mapping scheme. The pupil dilation features *P_d_* and *T* were quantified for each stimulation on/off epoch with awake mouse fMRI, enabling a PLR-based fMRI regression analysis. **Fig 5** showed the *P_d_*-based fMRI correlation patterns in both WT and AD mice. In both groups, there was a negative correlation observed in subcortical midbrain regions. For the WT mice, the correlation was in the serotonergic raphe nuclei, while the AD mice displayed this correlation predominantly in sparse pontine areas adjacent to the locus coeruleus (LC), laterodorsal tegmental area (LDT), and the mesencephalic pedunculopontine tegmental nuclei. Also, *P_d_*-based negative correlation with the cingulate cortex, hippocampus, and lateral septal (LS) area were detected in AD mice. Upon the group differential analysis, the most salient differences were detected in the MnR, the cingulate/retrosplenial cortex, the hippocampus (dentate gyrus (DG) and CA1), and the LS of the basal forebrain regions.

**Figure 5.**
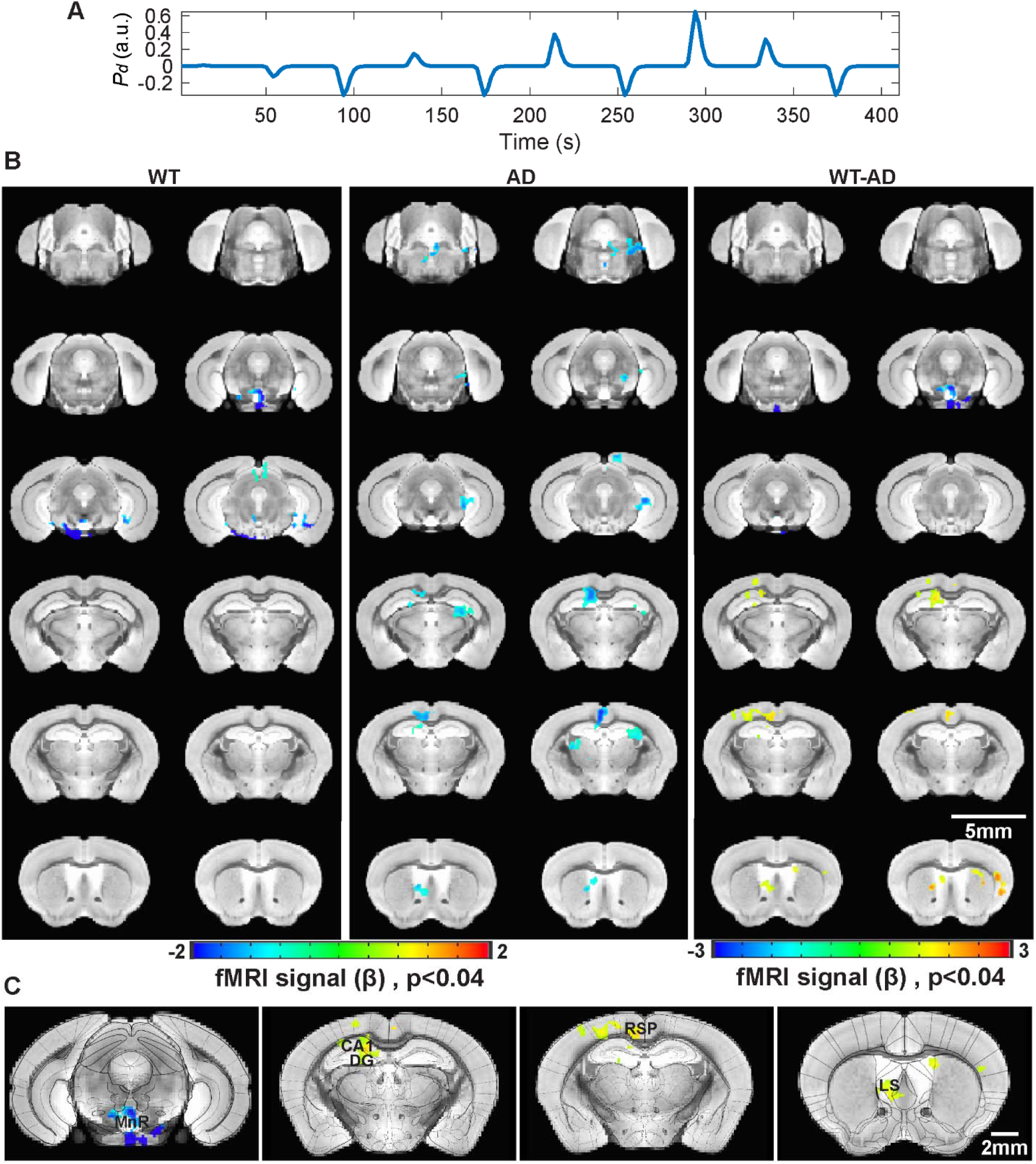
Differential analysis of *P_d_*-based fMRI correlation maps between WT and AD mice. A) A representative regression function based on epoch-specific *P_d_* features. The normalized positive/negative values indicate the deviation from the mean *P_d_* calculated from the given trial. B) The averaged brain-wide *P_d_*-based correlation maps of WT and AD mice (upper panel: WT, n=9 mice with 20 sections; middle panel: AD, n=9 mice with 19 sections), and the differential maps between WT and AD mice (lower panel: WT-AD, p<0.04, minimum cluster size=150 voxels, FDR correction=99.92%). C) The differential maps overlaid with the brain atlas highlight brain regions with significant difference: midbrain raphe nuclei (MnR), hippocampus (CA1, DG), retrosplenial cortex (RSP), and lateral septal nucleus (LS).

**Fig 6** showed the *T*-based fMRI correlation patterns, highlighting the negative correlation in the PRN, the cingulate cortex, and the hippocampus in the WT mice. There was also negative correlation scattered along the midbrain reticular formation and central thalamic nuclei in WT mice. Here, the AD mice showed positive correlation across the hippocampus and cingulate cortex, in addition to the hypothalamus. The statistical differential maps between WT and AD mice highlighted the most significantly different brain regions located at the PRN, the hippocampus, the cingulate/retrosplenial cortex, and central thalamic nuclei including the paraventricular (PVP) and dorsal thalamus.

**Figure 6.**
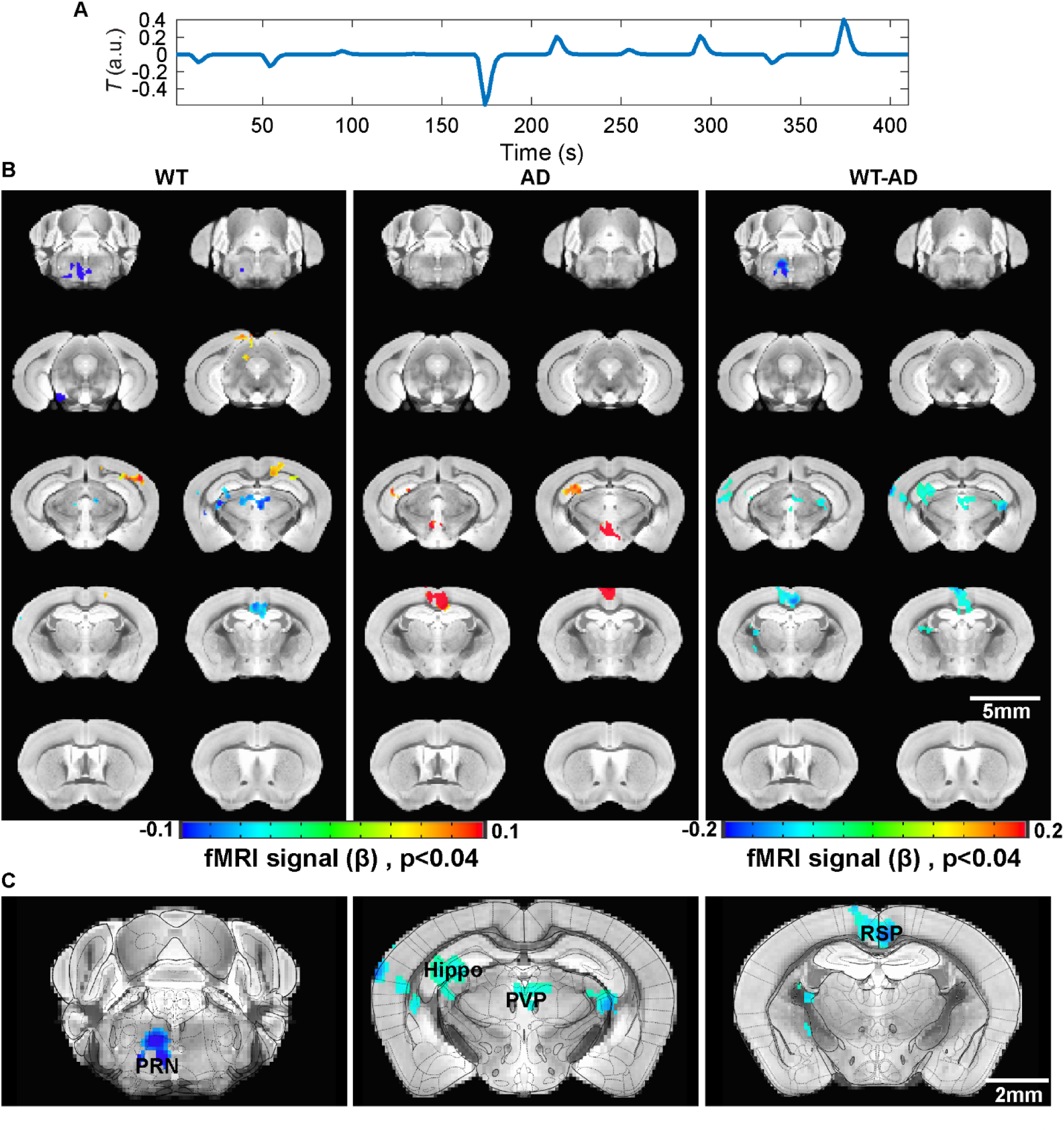
Differential analysis of *T*-based fMRI correlation maps between WT and AD mice. A) A representative regression function based on epoch-specific *T* features. The normalized positive/negative values indicate the deviation from the mean *T* calculated from the given trial. B) The averaged brain-wide *P_d_*-based correlation maps of WT and AD mice (upper panel: WT, n=9 mice with 20 sections; middle panel: AD, n=9 mice with 19 sections), and the differential maps between WT and AD mice (lower panel: WT-AD, p<0.04, minimum cluster size=200 voxels, FDR correction=98.35%). C) The differential maps overlaid with the brain atlas highlight brain regions with significant difference: pontine reticular nucleus (PRN), hippocampus (CA1, DG), paraventricular (PVP), and retrosplenial cortex (RSP).

To further identify the neuromodulatory nuclei responsible for the altered pupil-fMRI coupling in AD brains, we also performed ROI-based correlation network analysis given the reiterated visual stimulation paradigms. **Fig. 7A** illustrated the spatial correlation network of functional connectivity among the highlighted nuclei from PLR-based differential maps between WT and AD mice. By comparing the ROI-based correlation between WT and AD mice, significantly altered connectivity among paired functional nuclei was highlighted in the spatial correlation network maps, showing increased connectivity between hippocampus-PRN, LS-PRN and decreased LS-VC/SC, ACA-VC/SC in AD mice. **Fig 7B** showed the corresponding correlation matrices, summarizing the paired functional nuclei with significantly different connectivity between the two groups. It should be noted that we also performed permutation controls of randomly selected brain-wide ROIs for differential analysis, showing a dramatically decreased number of different connections in contrast to the selected nuclei based on the PLR-based fMRI analysis (**Fig 7C**). Meanwhile, brain-wide correlation analysis of ROIs, independent of the PLR-based fMRI mapping results, was performed to extract the ROIs with most significantly altered connectivity in AD mice. The top 20 paired ROIs with significantly altered connectivity between WT and AD mice, highlighted over 40% contribution to the connectivity changes originating from PRNc/PRNr in AD mice (**Fig 7D**), which was identified by the PLR-based fMRI differential analysis. These results further confirmed that the PLR-based fMRI differential analysis identified a critical list of neuromodulatory nuclei most responsible for AD-specific PLR.

**Figure 7.**
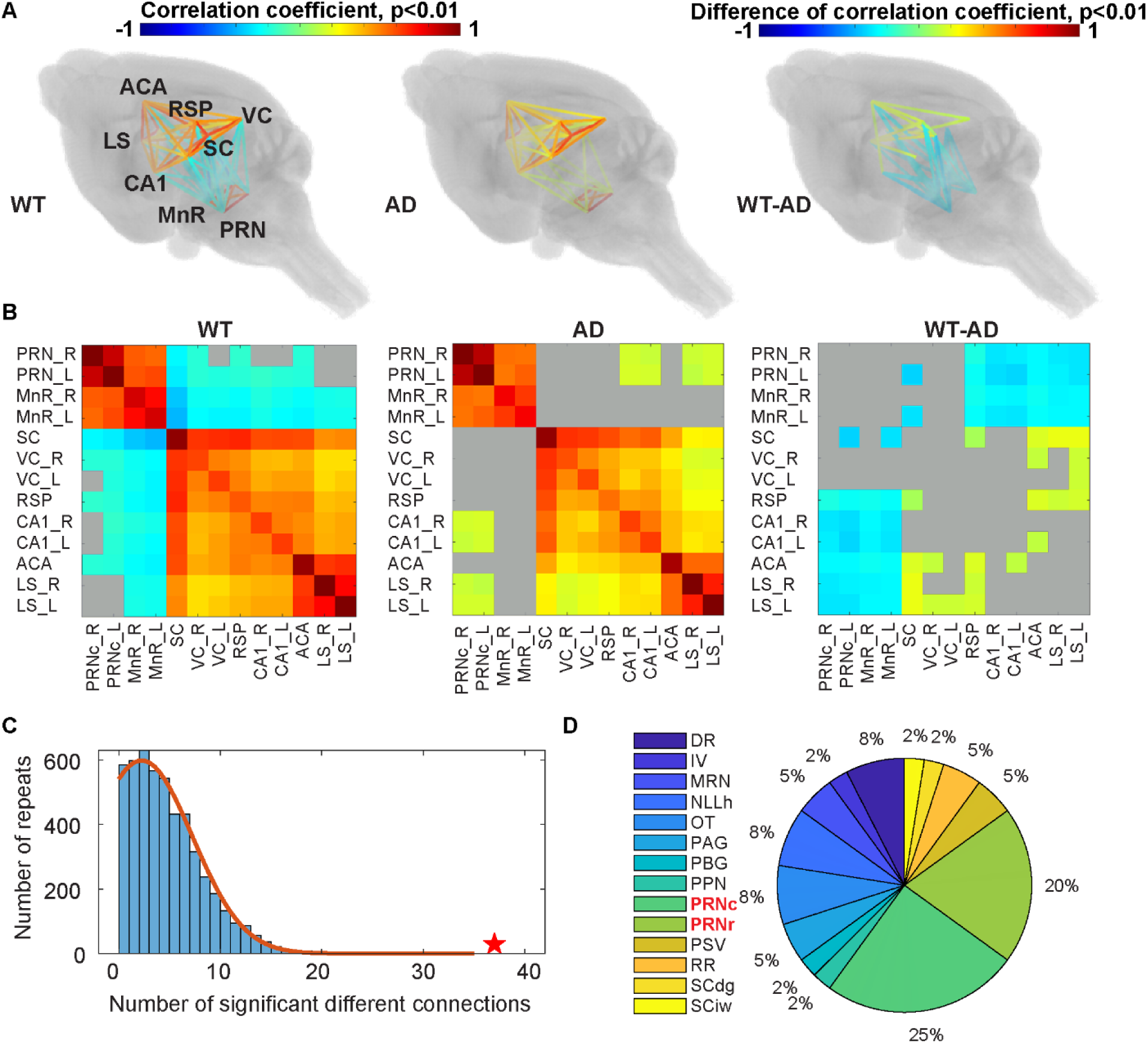
ROI-based correlation network analysis. A) Colored-coded Pearson correlation network through line projections in 3D mouse brain contours, which is based on the ROIs defined by PLR-based fMRI analysis (PRN, MnR, SC, VC, RSP, CA1, ACA, and LS). The color-coded line projections present the paired ROIs with statistically stronger based on one-sample t-test of WT and AD mice (WT: n=9 mice with 21 sections, AD: n=9 mice with 21 sections, p<0.01), and with significantly different correlation coefficients between WT and AD mice (WT-AD, two-sample t-test, p<0.01), B) The correlation matrices of WT and AD mice, as well as the differential matrix (WT-AD), presenting an alternative way to show the correlation networks based on the PLR-fMRI specific ROIs. C) The permutation test histogram shows the distribution of number of significantly different connections between WT and AD mice across 5000 permutation tests with randomly selected ROIs, which was fitted with a Gaussian distribution profile (red line, μ=2.25, σ=5). Red star indicates the number of significantly different correlation based on the PLF-fMRI specific ROIS, which is ∼μ+7σ of the Gaussian distribution. D) A pie chart highlights the ROIs presenting the most different correlation patterns (i.e. the top 20 paired ROIs with largest coefficient difference at p<0.01) between WT and AD mice, of which the PRN accounts for 45%)

## Discussion

The present study combined brain-wide awake mouse fMRI with real-time pupillometry to elucidate the linkage of neuromodulatory function and AD-specific pupil dynamics. This preclinical pupil-fMRI mapping scheme in awake mice provides a proof-of-concept method to identify non-invasive pupillary biomarkers coupled with altered neuromodulatory function due to AD brain degeneration.

PLR in AD patients has been reported to show altered pupil constriction phases ^24–26^. These studies have suggested the potential of pupil dynamics as biomarkers of AD degeneration. However, a few studies fail to reproduce altered PLR features when comparing AD patients with control groups^27,28^. Numerous efforts have been made to bridge pupil dynamics with diversified behavioral variables such as subject’s arousal fluctuation ^5–7^, designated tasks ^5,8–12^, or specific cognitive processes ^13–21^. The richness of behavioral correlates of pupil dynamics, in particular, for AD patients with cognitive impairments, would lead to confounding PLR given the complex controlling mechanisms. The present study analyzed PLR in awake AD mice, showing illumination-evoked pupil constriction phase with reduced amplitude of pupillary constriction (*P_c_*) (p=0.0383) among 580 visual stimulation epochs of 9 WT mice and 600 visual stimulation epochs of 9 AD mice In contrast to the pupillary constriction alteration, the post-illumination pupillary recovery phase showed a significant difference in AD mice, demonstrating shorter recovery time (T) (p=3e^-10^) and reduced pupil dilation amplitude (P) (p=0.001). Despite the reported statistical significance, the PLR in awake mice showed large variance across different stimulation epochs, confounding the PLR as biomarkers of AD brains for case-specific characterization. The phasic regulation of PLR relies on both sympathetic and parasympathetic systems innervating muscles controlling pupil dilation and constriction^29–31^. Beside the brainstem reflective function underlying PLR^67^, the higher level functional demand of visual perception and attention is well incorporated through the pupil dynamic controlling mechanism^68–70^. Furthermore, the activity of subcortical nuclei mediating neuromodulation has been tightly coupled with pupillary responses^33,43,51,56,57,71,72^. These studies well supported the present work to extract the event-related PLR phase-dependent fMRI correlation across different trials.

One intriguing technological development of this work is the high-resolution awake mouse fMRI at 14T. Hike et al developed a novel implantable RF coil, which can also be used as the headpost for head-fixation during awake mouse fMRI scanning ^64^. In contrast to the inductive coupled wireless RF coil ^73,74^, this new design has enabled ultra-high-resolution EPI-based fMRI of awake mice with 100×100×200um resolution, permitting the characterization of activated functional nuclei in mice only a few hundred micron size. Both AD and WT mice have gone through the acclimatization procedure with fMRI as describe previously ^75^, enabling the reliable detection of visual stimulation-evoked fMRI maps (**Figs. 2 & 3**). It should be noted that the functional maps showed brain-wide activation not only at the visual pathway, but also included the thalamic areas and ACA in both groups. This demonstrated extended high level cortical function involved in cognitive processing of awake mice. This advantageous awake mouse fMRI allowed the brain-wide functional characterization of AD-specific PLR features.

We have performed PLR-based fMRI correlation analysis based on the stimulation on/off epoch across different trials. In contrast to the direct comparison of evoked functional maps between WT and AD groups, the *P_c_*-based AM regression analysis, highlighted a critical modulatory effect based on the illumination-driven retinal activity. The BOLD signal difference detected along the central visual pathways indicated significantly reduced visual cortical activity from primary and associated cortices in AD mice, as well as in the LGN, but no signal difference was detected in the central thalamic areas and cingulate cortex (**Fig 3A**). This altered visual pathway activity in AD mice could be directly linked with the impaired retina function due to vascular and neuronal degenerative effect ^76–79^. The differential maps highlighted V1 and associated visual areas with LGN, indicating impaired high-order visual signal processing. Previous studies have reported altered laminar distribution of neurofibrillary tangles (NFTs) in the visual cortex ^80^ and decreased long synaptic projection from area V2 to V1 in AD patients ^81^.

The *P_c_*-based fMRI provides a bottom-up mapping scheme to characterize the malfunction of visual signal processing in AD brains. In contrast to the *P_c_*-based fMRI correlation, the post-illumination pupil dilation reflected the top-down neuromodulatory processes, of which *P_d_*/T-based fMRI revealed altered coupling with multiple s functional nuclei in AD mice. First, the posterior cingulate cortex and hippocampus of AD mice showed altered coupling features with *P_d_*/T when compared to WT mice. Since both the posterior cingulate cortex and hippocampus are impaired in AD due to the neurodegeneration ^82–86^, it is plausible that the reduced neuronal activation in these areas was directly involved in the altered pupil dilation features, especially the shortened dilation recovery time that presented the most significant changes in PLR measurements (**Fig 1C**). This observation is also consistent with the reports, showing that damage to the hippocampus in AD patients is linked with the impaired oculomotor behavior and networks^87–89^.

Another intriguing observation is the identification of subcortical brain regions underlying the altered pupil dilation recovery in AD mice. First, the PLR (T)-based fMRI coupling in PRN was reduced in AD mice compared to WT mice. The PRN is the major relay nuclei in the brainstem controlling oculomotor function^90–92^. By performing the brain-wide ROI network connectivity analysis based on the reiterated visual stimulation (**Fig 7D**), significant changes in PNR-based network connectivity in AD mice further confirmed the PLR (T)-based PRN coupling as a crucial estimate of the reliable pupil-fMRI biomarker for AD brains.

For the *P_d_*-based fMRI correlation, negative correlation with median raphe nuclei (MnR) was shown in WT mice and less coupled in AD brains, indicating impaired serotonergic regulation of pupil dilation. The pupil dilation and constriction can be controlled by serotonergic agonists and antagonists, respectively ^93–95^, and significant neuronal reduction is also reported in the MnR of AD patients’ brains ^96^ In contrast, the LS showed more negative correlation in AD than WT mice in the *P_d_*-based fMRI maps. LS is composed predominantly of GABAergic neurons and is reciprocally connected to multiple subcortical regions^97^, in particular, the medial septal area (MS) that sends cholinergic projections throughout the cortex^98^. Cholinergic hypofunction has been reported in AD brains^99,100^, showing the impaired cholinergic projections to broad brain areas^101–107^. Since the cholinergic function is coupled with the pupil dilation directly^56^, the increased negative correlation in LS indicates that LS-mediated septal cholinergic function in AD brains contribute more to control the post-illumination pupillary dilation than healthy brains.

A counter-intuitive observation in this work is the primary negative correlation patterns from multiple nuclei for the *P_d_*-based fMRI maps. The pupil-based fMRI negative correlation has been previously reported in anesthetized rodent brains ^51^. And brain-wide negative BOLD signal was also coupled the vigilant brain state changes based on eye open/close arousal indices ^50^. The P-based negative BOLD fMRI correlation could be mediated by the neurovascular coupling event underlying arousal state changes in awake mice involving multiple neuromodulatory pathways. By incorporating the resting-state spatiotemporal pupil-fMRI correlation patterns, principal component analysis (PCA) has been implemented to highlight different neuromodulatory pathways based on clustered pupil dynamic spectra ^108^. To further elucidate the neuronal basis of the negative coupled pupil-fMRI relationship, cellular-specific fiber photometry Ca^2+^ recordings ^51^ need to be implemented with awake mouse fMRI and real-time pupillometry.

In conclusion, we have shown that the epoch-specific post-illumination PLR features, i.e. the pupil dilation recovery time (*T*) and amplitude (*P_d_*), could be correlated with specific neuromodulatory nuclei that can exhibit impaired functionality in AD brains. These multiple nuclei also represented dramatically altered network connectivity during visual stimulation in AD mice (**Fig 7**), further indicating the AD-specific PLR features with underlying altered neuromodulatory function. This pupil-fMRI mapping scheme could provide non-invasive measurements to identify pupillary biomarkers of brain degeneration in AD patients.

## Materials and Methods

### Animal surgical procedures

Thirteen C57BL/6J WT mice and Nine 5xFAD transgenic AD mice were used in the present study (weighing ∼20 to 30 g). For the brain-wide fMRI datasets, thirteen WT mice were used for data analysis. For real-time pupillometry with fMRI, only nice of the thirteen WT mice were used because of the poor quality of the pupil recorded inside the 14T MRI scanner. Mice were group housed (2-4/cage) under a 12-h light/dark cycle with food and water ad libitum. All animal procedures were conducted in accordance with protocols approved by the Massachusetts General Hospital (MGH) Institutional Animal Care and Use Committee (IACUC), and animals were cared for according to the requirements of the National Research Council’s Guide for the Care and Use of Laboratory Animals.

All of the mice underwent surgery under anesthesia (1-2% isoflurane and 1L/min of medical air and 0.2L/min additional O^2^ flow) to implant the RF coil to the skull as described in our previous study ^64^. Briefly, anesthetized mouse heads were stabilized in a stereotaxic stage. And the skull was exposed after the aseptic treatment. Before the coil was implanted, both 0.3% H_2_O_2_ and PBS was applied to clean the skull. It is critical to wait until the skull is fully dried before the coil implantation. The coil was secured with cyanoacrylate glue and dental cement to prevent movement and ensure stability. Following surgery, mice received medications, and were allowed at least a week of recovery in their home cages for neck strengthening and to resume normal head movements.

Animals were acclimated to the MRI environment through a modified training program spanning five consecutive weeks, progressively increasing from 10 minutes in Week 1 to 60 minutes in Week 5, following established procedures adapted from prior research ^75^.

#### MRI methods

MRI data was acquired with the 14T horizontal MRI scanner (Magnex Sci, UK) located at the Athinoula A. Martinos Center for Biomedical Imaging in Boston, MA. The magnet is equipped with a Bruker Avance Neo Console (Bruker-Biospin, Billerica, MA) and is operated using ParaVision 360 V.3.3. A microimaging gradient system (Resonance Research, Inc., Billerica, MA) provides a peak gradient strength of 1.2T/m over a 60-mm diameter. The MRI data for anatomical registration was obtained with a multi-slice T1-weighted 2D gradient echo Fast Low Angle SHot (FLASH) sequence (TE=3ms, TR=475ms, flip angle = 30°, 4 averages for an approximate acquisition time of 4.5 minutes, 100×100×200µm resolution). The awake mouse fMRI images were obtained with a multi-slice 2D EPI (TE: 6.2ms, TR: 1s with two segments, BW: 277,777Hz; Matrix size: 144×96×36, in-plane resolution: 100×100µm, thickness: 200µm). A total of 205 repetition is acquired per trial for a total of 6 min 50s. For each mouse, three trials of visual stimulation were conducted per scanning session. Within each trial, a block-design paradigm consisting of 10 stimulation on/off epochs was applied. For each epoch, the visual stimulation lasted for 8 seconds, followed by 32 seconds of inter-stimulus intervals. The optical stimulation during the “8s on” period included two LED lights flashing at specific frequencies: 490nm at 5.1Hz and 530nm at 5Hz, with light pulse duration at 20 ms. Before the stimulation, five scans were obtained as a baseline.

### Pupillometry setup during fMRI scanning

The pupil recordings were obtained using a 3D-printed animal cradle with mounts for a camera, a mirror, and three fiber optic cables used as light sources as shown in **Fig 1A**. The pupil dynamics were recorded through the mirror. The red light (660nm) was used for video illumination since mice are not sensitive to wavelengths over 600nm^63^, while the green and blue lights (530nm and 490nm) were used for visual stimulation.

To correlate the whole-brain fMRI signals with specific pupillary response features of normal and AD mice, detection of pupil dynamics is first needed. **Fig 1B** shows an example of pupil dynamic recording. As shown in **Fig 1A, t**he edge of the mouse pupil was detected and tracked using DeepLabCut ^65^. To further confirm the PLR results, an alternative pupil size detection method based on ocular and global grayscales was proposed. By comparing the average grayscale of the eye with the average grayscale of the whole image, the size of the pupil could be well characterized based on the signal intensity (**Supplementary Fig 1**). Eventually, the normalized pupil dynamics data *P* can be calculated as:

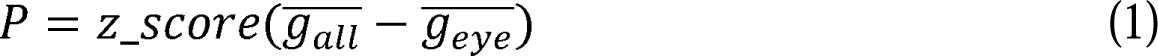

Where *g_eye_* is the grayscale of the eye area and *g_all_* is the grayscale of the whole image.

### PLR features extraction

To further map the correlation between pupil and fMRI, we extracted different features of the PLR in AD and WT mice. The PLR time courses were divided into two components (constriction and dilation) for phase-dependent analysis of PLR. The constriction component represents the immediate reaction of the pupil to illumination, primarily regulated by the parasympathetic nervous system. To quantify pupil constriction, we utilized the difference of the trial-specific z-scored pupil dynamic data before and after constriction, denoted as *P_c_*. For each epoch, the pre-constriction pupil size is calculated by averaging the pupil dynamic recording 1 second before the stimulation. The constricted pupil size is calculated by averaging the pupil recording below 75% of the epoch-specific minimum z-score. The dilation component, mainly controlled by the sympathetic nervous system, represents automatic pupil modulation, and can provide insights into associated brain activity. Regarding the dilation component, we used an exponential function to accurately model the changes in pupil size.

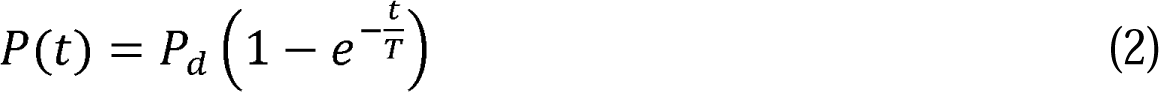

Where P(t) is the pupil dilation defined from above 75% of the epoch-specific minimum z-score after stimulation to 75% of the epoch-specific maximum z-score. Through this modeling approach, we obtained two parameters: *P_d_*, indicating the difference in pupil size before and after dilation; *T*, characterizing the shape of the pupil dilation curve and the potential full dilation time.

### BOLD fMRI mapping with visual stimulation

To generate the visual-stimulation BOLD map, the fMRI images were processed using Analysis of Functional Neuroimages (AFNI) software^109,110^, i.e. the “anfi_proc.py”. For group analysis, we identified the mouse brain template at 100µm isotropic resolution, which is resampled from the high-resolution images acquired from Australian Mouse Brain Mapping Consortium (AMBMC)^111^. For each experiment, the anatomical FLASH image was registered to the AMBMC template using 3dAllineate function and the transformation matrix was created. Meanwhile, averaged EPI data was registered to the anatomical image and corresponding transformation matrix was created based on the registration. By applying the two transformation matrices, EPI time courses from multiple trials per experiment will be registered to the brain template for concatenation. In addition, outlier of the time courses was characterized and a censoring function was applied to remove the erroneous data points showing large image distortion due to motion as detected by outlier function. Several pre-processing procedures were implemented before the general linear regression-based denoising, such as despike, volreg, mask, blur, and scale to ensure compatibility and consistency across the analysis pipeline. 3dDeconvolve function was used to create the final statistical β coefficient maps using a block function. A detailed awake mouse fMRI imaging processing procedure has been described previously ^64^.

### PLR feature-driven fMRI map

To characterize PLR feature-driven fMRI maps, we applied the stimulation on/off epoch-specific PLR features to modulate the regression function. For group analysis, voxel-wise Student’s t-tests were performed to identify the differential PLR-based fMRI correlation maps between AD and WT mice.

To analyze the constriction component, pupil constriction parameter *P_c_* was applied to the amplitude-modulated (AM) pupil-fMRI correlation analysis. The *P_c_* was used as the auxiliary behavioral information to modulate the amplitude of hemodynamic response function. The AM1 function of AFNI was applied with a regressor calculated as the following:

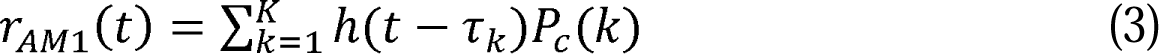

Where 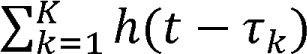 is the ideal HRF, *P_c_* (k) is the value of the *P_c_* at time *k*. *K* is the time length of the experiment trail.

For the dilation component, pupil dilation parameters *P_d_* and *T* were used as the auxiliary behavioral information which was expected to be modulated by activation in specific brain areas. Those brain areas were expected to be associated with automatic pupil modulation and to exhibit variation between AD and WT groups. The AM2 function of AFNI was applied to incorporate the ideal HRF, as well as two regressors presenting the epoch-specific modulation for pupil dilation parameters (*P_d_* and *T*), which were calculated as the following:

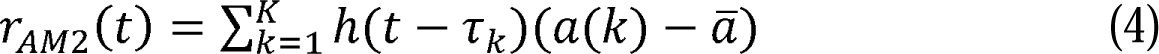

Where a(k) is the value of the pupil dilation parameter at time *k*. a is the average pupil dilation parameter.

For the statistical differential maps, significant voxels were presented based on the p-values less than 0.04 to 0.001 according to different comparison schemes. The false discovery rate correction was found to be over 98%. The minimal cluster size was set at 150 or 200 voxels according to different comparison schemes.

### Functional network connectivity analysis

For the correlation network analysis, we defined all the ROIs based on the Allen mouse brain atlas given the brain-wide delineation of the functional nuclei. This Allen mouse brain atlas was registered to the previously chosen AMBMC brain template. For each ROI, we averaged the time course from the preprocessed fMRI datasets based on the 21 sections of 9 WT mice and 21 sections of 9 AD mice. Pearson correlation analysis was performed to calculate the correlation coefficients among paired ROIs that were identified based on the PLR-feature-driven BOLD differential maps between WT and AD mice (a total of 13 ROIs were selected). One-sample Student’s t-test was performed to identify the significantly correlated paired ROIs in two groups of animals. Two-sample Student’s t-test was performed to compare the correlation difference between the paired ROIs of the two groups. To verify the differential correlation networks of PLR-specific ROIs, permutation control tests were performed by randomly selecting same number of ROIs (excluding those pupil feature-related ROIs) for 5000 times. For each permutation test, the number of randomly paired ROIs with significantly different correlation between two groups was calculated. Gaussian fitting was applied to verify the normal distribution for the permutation tests. Alternatively, Pearson correlation analysis was also performed across all the ROIs based on the Allen mouse brain atlas (a total of 1086 ROIs) and the two-sample Student’s t-tests were performed to compare correlation coefficients of paired ROIs between two groups. Among the ROIs showing the significant correlation difference between two groups (p<0.01), the top 20 paired ROIs with largest coefficient difference were chosen to plot the pie chart, which shows the individual ROIs’ contribution to the altered correlation network between WT and AD mice. **Supplementary Fig 3** also showed the pie charts from top 10 to top 200 verified paired ROIs.

## Author Contributions

XL and XY designed the research. XL, DH, YJ, and XY performed the research. XL, SC, and WM analyzed data. CR provides animal models. XAZ performed surgeries. DH, XY built coils. DH and XL designed the animal cradle. XL, DH, and XY wrote the paper.

## Acknowledgements

We thank Zeping Xie, Bei Zhang, and Biyue Zhu for their technical support. The research is supported by funding sources, including NIH grants (RF1NS113278, RF1NS124778, R01NS122904, R01NS120594, and R21NS121642), U19 Cooperative Agreement Grant (U19NS123717), S10 instrument grants (S10OD028616 and S10RR025563) to Martinos Center, Massachusetts General Hospital, and NSF grant (2123970).

