## Supplemental information for "Mapping the bioimaging marker of Alzheimer’s disease based on pupillary light response-driven brain-wide fMRI in awake mice"

**Author Contributions:**

XL and XY designed the research. XL, DH, YJ, and XY performed the research. XL, SC, and WM analyzed data. CR provides animal models. XAZ performed surgeries. DH, XY built coils. DH and XL designed the animal cradle. XL, DH, and XY wrote the paper.

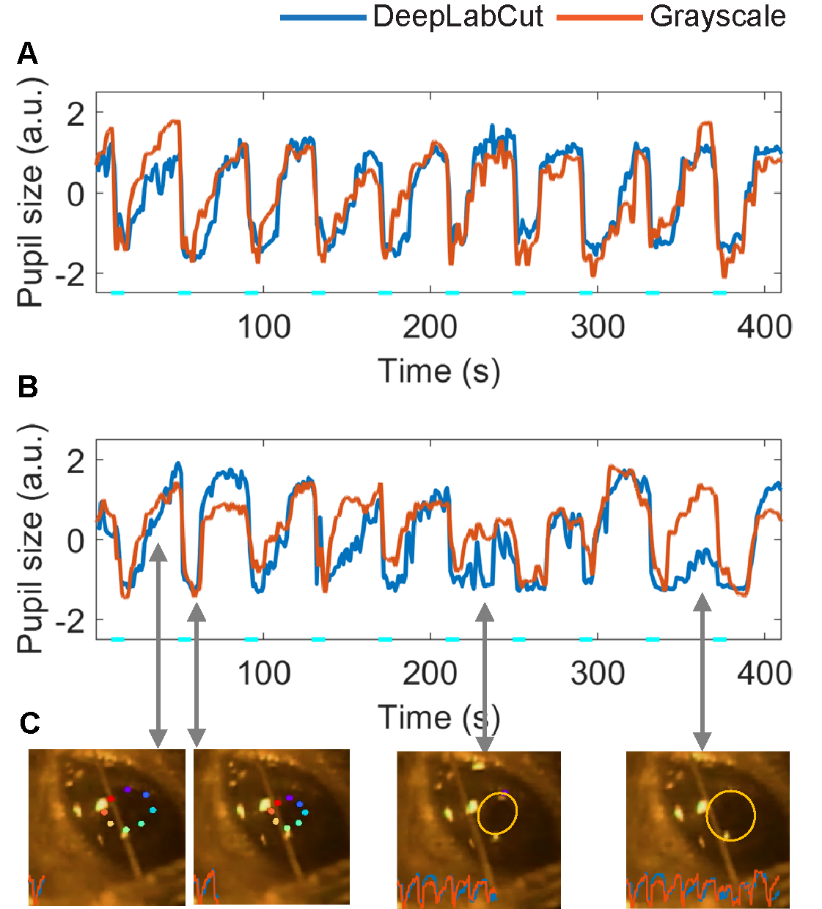

**Supp Figure 1**. **PLR measurements based on the grayscale-based method and DeepLabCut.** A) The representative plots showing the similar detection capability of pupil size changes by grayscale-based method and DeepLabCut. B) The representative plots showing the deviation of pupil size detection by the grayscale-based method in comparison with DeepLabCut due to external confounding factors (i.e. whisker motion in front of the eye). C) The corresponding pupil recording frames used for pupil size detection in B (A whisker can be seen interfering with pupil measurements).

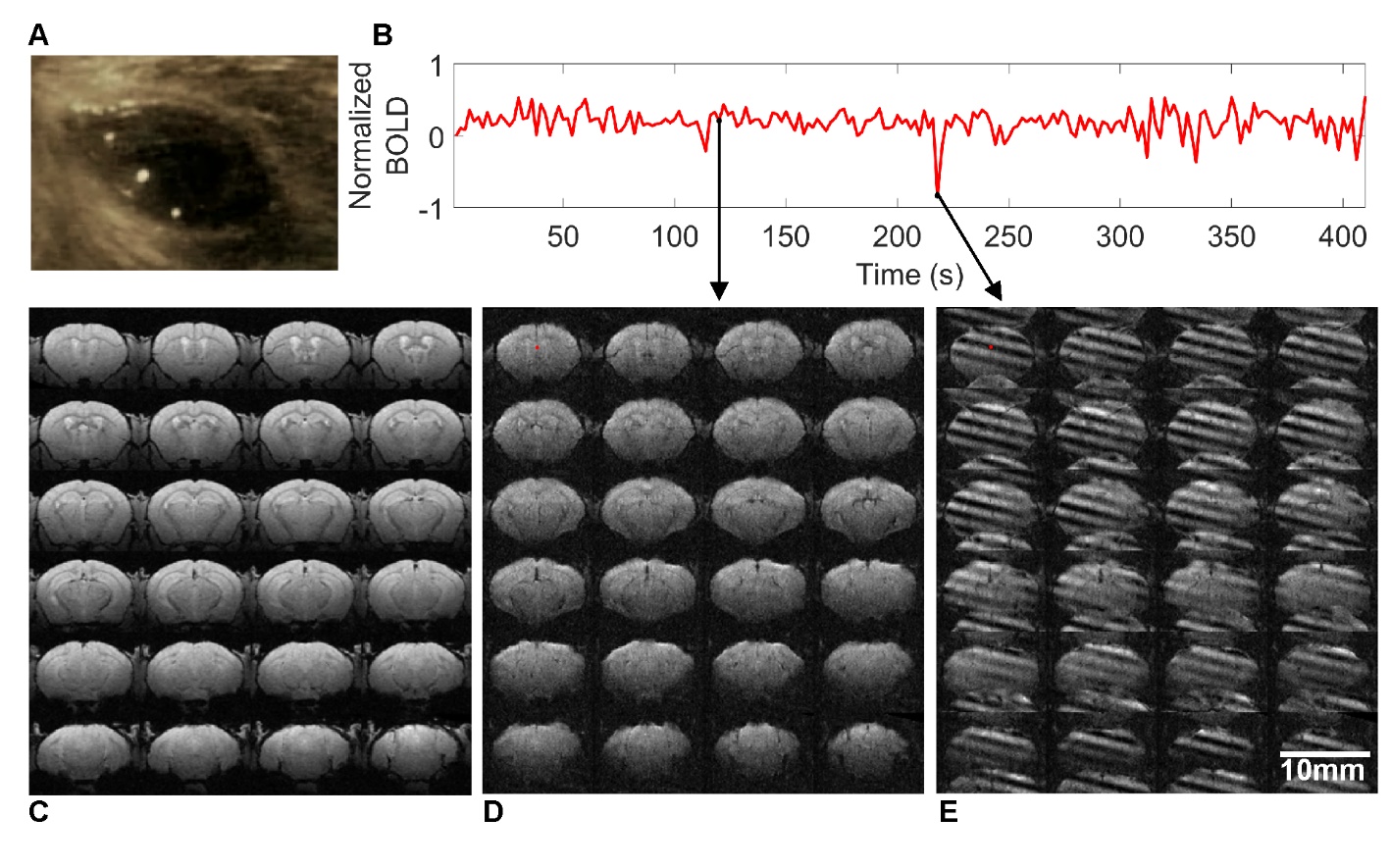

**Supp Figure 2. High-resolution awake mouse fMRI with real-time pupil dynamics recording at 14T.** A) A frame of the pupil recording of awake mice during fMRI scanning inside the 14T scanner. B) The representative fMRI time course from the selected voxel of the high-resolution raw EPI images, enabling the trace of motion-induced artifacts. C) The brain-wide anatomical MRI images (FLASH) with minimal susceptibility. D) The snapshot of the raw EPI fMRI image without motion artifacts. E) The snapshot of the distorted images due to the motion of the awake mouse during scanning (**Supplementary Movie 1** shows the video of motion-induced artifacts throughout the fMRI trial).

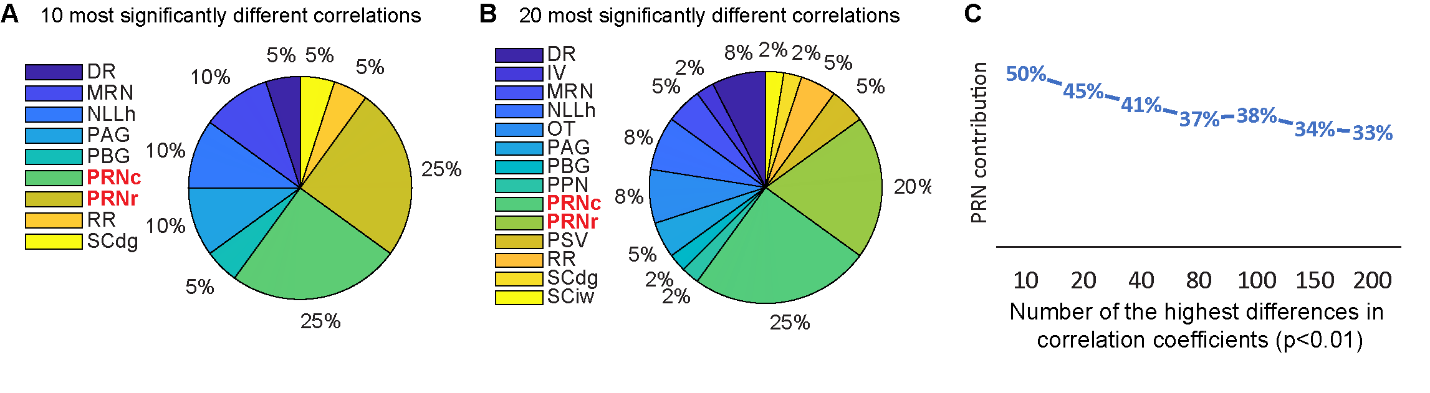

**Supp Figure 3 The strongest significant differences between WT and AD mice.** A) A pie chart of 10 strongest differences of correlation coefficient between WT and AD mice, of which PRN accounts for 50%. (two-sample t-test, p<0.01). B) A pie chart of 20 strongest differences of correlation coefficient between WT and AD mice, of which PRN accounts for 45%. (two-sample t-test, p<0.01.) C) Graph showing level of PRN contribution to the highest significantly different correlation coefficients emphasizing the important role of PRN. (two-sample t-test, p<0.01)

**Supp Movie 1**. **Real-time awake mouse EPI images and pupil dynamics recording during visual stimulation.** The video showed real-time EPI raw images and pupil dynamics recording from the awake mouse during visual stimulation. The real-time tracer from the selected voxel, which is paired with the raw EPI images, includes the time points with motion-induced image distortion during awake mouse fMRI.
